# PPa1 insufficiency drives lysosomal storage disease and inflammatory macrophage expansion in the bone marrow

**DOI:** 10.64898/2026.03.16.712247

**Authors:** Magdalena Grzemska, Luming Chen, Jamie Russell, Nagesh Peddada, Samantha Calvache, Jianhui Wang, Aysha Khalid, Jonathan J Rios, Jeff SoRelle, Bruce Beutler, Evan Nair-Gill

## Abstract

Inorganic pyrophosphatase-1 (PPa1) is an essential enzyme proposed to limit accumulation of the ubiquitous metabolic byproduct inorganic pyrophosphate, yet its role beyond a general housekeeping function remains poorly understood. We generated viable hypomorphic alleles of *PPa1* through random mutagenesis in mice and unexpectedly found that *PPa1* insufficiency causes a lysosomal storage disease of the bone marrow. Mutant mice developed impaired hematopoiesis and defective skeletal mineralization in a hematopoietic-intrinsic manner. The bone marrow was infiltrated by glycolipid-laden macrophages that were marked by high Spp1 and CD14 expression, blunted lysosomal acidification, metabolic stress, and a distinct inflammatory transcriptional program. Consistent with a lysosomal storage defect, *PPa1*-deficient bone marrow accumulated elevated levels of long-chain glucosyl-sphingolipids and sphingosine. These findings identify PPa1 as a previously unrecognized regulator of lysosomal function in myeloid cells that restrains inflammatory macrophage expansion and prevents bone marrow lysosomal storage pathology.

## INTRODUCTION

Intracellular soluble pyrophosphatases (sPPases) are highly conserved enzymes that hydrolyze inorganic pyrophosphate (PPi) into inorganic phosphate (Pi)^1, 2^. PPi is generated as a byproduct of most biosynthetic reactions, and its hydrolysis is thought to remove an energetically unfavorable reaction product, thereby driving the elongation of polymeric macromolecules^3^. Consistent with this central metabolic role, loss of sPPase activity leads to profound defects in cellular proliferation and viability^4–6^.

Despite their essential function, the roles of sPPases in maintaining cellular and organismal homeostasis remain poorly defined, in part because mammalian models of systemic sPPase deficiency have been lacking. Humans and mice express two sPPases with different subcellular locations: inorganic pyrophosphatase-1 (PPa1), which localizes to the cytoplasm, and inorganic pyrophosphatase-2 (PPa2), which localizes to mitochondria^6^. Biallelic mutations in *PPA2* cause sudden cardiac death and myocardial scarring through poorly understood mechanisms^7^. In mice, adipocyte-specific deletion of *PPa1* results in severe lipodystrophy and insulin resistance, indicating a tissue-restricted requirement for PPa1 in metabolic homeostasis^8^. More recently, biallelic *PPA1* mutations associated with reduced pyrophosphatase activity were identified in pediatric siblings with neonatal galactosemia^9^. However, no additional major clinical phenotypes were reported. The prevalence of deleterious biallelic *PPA1* variants in the human population, as well as their long-term clinical consequences, remain unknown.

Here, through forward genetic screening in mice, we discovered viable hypomorphic alleles of *PPa1*. Unexpectedly, homozygous mutant mice exhibited a very specific syndromic disease process. They developed severe defects in hematopoiesis and skeletal homeostasis which were accompanied by expansion of inflammatory macrophage populations and sphingolipidosis within the bone marrow, producing a phenotype that closely resembled classical lysosomal storage disorders (LSDs). Our findings establish a previously unrecognized link between PPa1, myeloid inflammatory programming, and lysosome function, suggesting that PPa1 may be involved in a broader spectrum of hematologic and metabolic diseases.

## RESULTS

### Random germline mutagenesis links a mutation in *PPa1* to defective hematopoiesis

To discover new determinants of hematopoiesis, we screened the peripheral blood of N-ethyl-N-nitrosourea (ENU) mutagenized mice with flow cytometry^10^. Several mice from a single pedigree showed decreased proportions of circulating neutrophils, a phenotype that we named *hotpot* (*hpt*). Automated meiotic mapping linked the *hpt* phenotype to two missense mutations in *inorganic pyrophosphatase-1* (*PPa1*) using a recessive model of inheritance (Fig. 1a, b).

**Figure 1:**
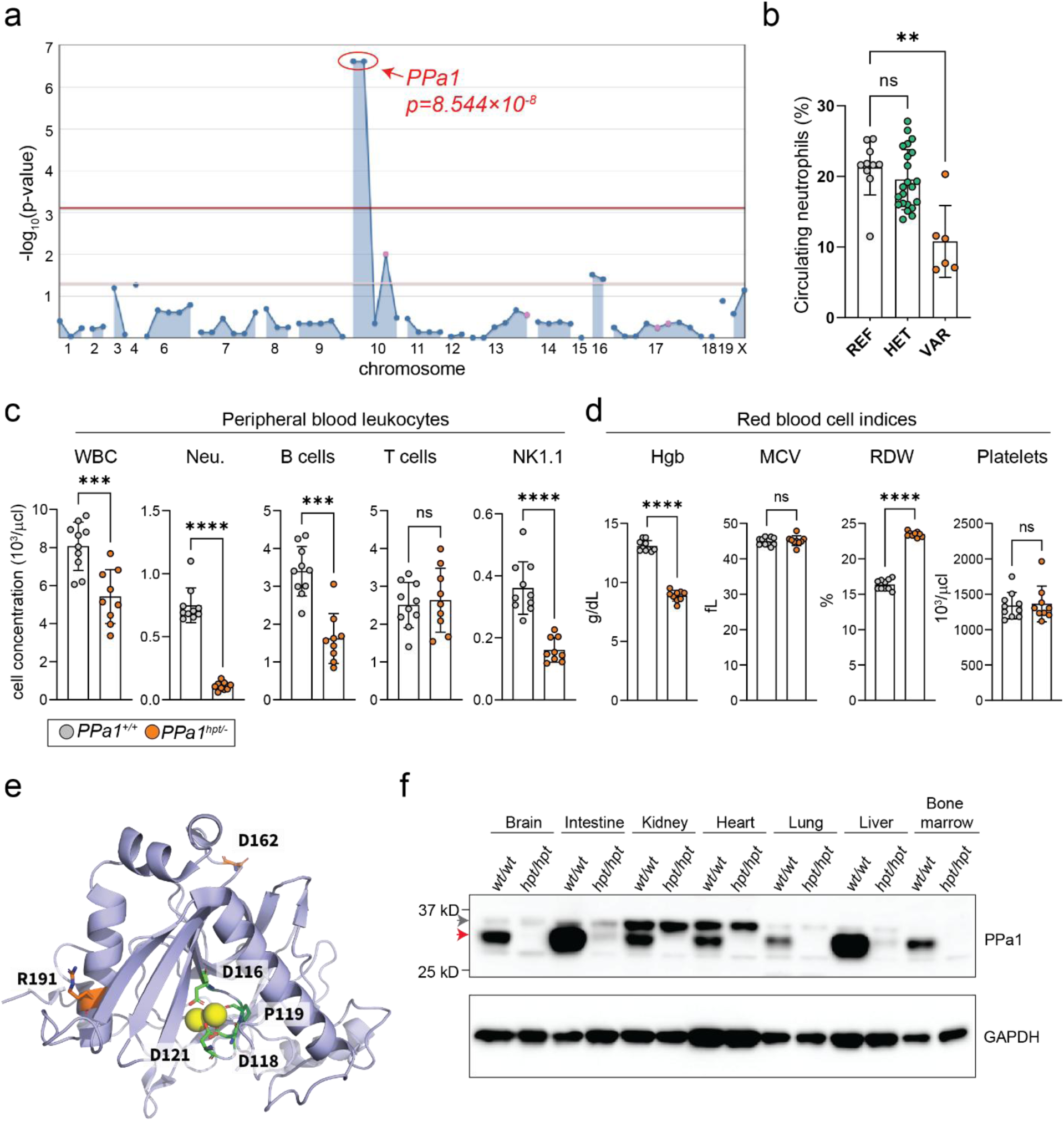
An ENU mutagenesis screen shows PPa1 to be important for peripheral blood cells counts. **(a)** Manhattan plot showing *P* values for linkage of two mutations in *PPa1* with the *hotpot* (*hpt*) neutrophil phenotype using a recessive model of inheritance. The −log_10_ *P* values (y-axis) are plotted versus the chromosomal positions of the mutations (x-axis). Horizontal red and purple lines represent thresholds of *P* = 0.05 with or without Bonferroni correction, respectively. **(b)** Frequency of circulating neutrophils plotted per genotype [REF *PPa1^+/+^* (n=10); HET *PPa1^+/hpt^*(n=22) ; VAR *PPa1^hpt/hpt^* (n=6)]. Each symbol represents an individual mouse. **(c)** WBC counts and **(d)** RBC indices in the peripheral blood of compound heterozygous mice carrying the *hotpot* allele and a CRISPR-induced knockout allele. Each symbol represents an individual mouse [*PPa1^+/+^* (n=10); *PPa1^hpt/-^* (n=9)]. **(e)** Crystal structure model of the human PPa1 monomer (PDB: 7BTN) with the active site residues (green), Mg^2+^ (yellow balls), and R191 and D162 (orange) highlighted. **(f)** Endogenous expression of PPa1 in various tissues in WT and *hotpot* mice. The red arrow indicates the expected band, and the gray arrow indicates a background band. Data are expressed as mean ± s.d. and significance was determined using the Kruskal-Wallis test with Dunn’s multiple comparisons **(b)** and the Mann-Whitney test **(c, d)**.

To confirm that the mutations in *PPa1* caused the *hpt* phenotype, we crossed the *hpt* strain to mice carrying a heterozygous *PPa1* knockout allele generated by CRISPR-Cas9 gene editing. Compound heterozygotes (*PPa1^hpt/-^*) displayed decreased circulating total white blood cells (WBCs) and neutrophils implicating *PPa1* as the causative gene of the *hpt* phenotype (Fig. 1c). *PPa1^hpt/-^* mice also showed decreased circulating B cells and NK1.1 natural killer cells but normal numbers of circulating T cells (Fig. 1c). In addition to WBC deficiencies, *PPa1^hpt/-^*mice were anemic (Fig. 1d). Red blood cells (RBCs) exhibited a normal mean corpuscular volume (MCV) with an elevated red cell distribution width (RDW) indicative of anisocytosis, a common finding during chronic inflammation and infiltrative bone marrow diseases^11^. The number of platelets was normal (Fig. 1d). Together these data show that recessive mutation of *PPa1* causes deficiencies in multiple blood cell lineages suggesting defective hematopoiesis.

The *hpt* allele encoded an arginine to glycine substitution at position 191 (R191G) and an aspartic acid to glycine substitution at position 162 (D162G). Based on the crystal structure of the highly conserved human PPa1 protein, both R191G and D162G were located away from the catalytic site (Fig. 1e)^12^. Using AlphaMissense, the R191G mutation was predicted to be damaging (pathogenicity score: 0.576), whereas the D162G mutation was predicted to be benign (pathogenicity score: 0.09)^13^. The effect of the *hpt* allele was to drastically reduce endogenous PPa1 protein expression across various tissues (Fig. 1f).

### PPa1-deficiency leads to hematopoietic and skeletal defects

*PPa1^hpt/-^* mice were born at a sub-Mendelian ratio (Fig. S1a). To routinely generate enough mice for experiments, we used *PPa1^hpt/hpt^* mice which were born at a Mendelian ratio while retaining the severe hematologic phenotypes found in compound heterozygotes (Fig. S1b, c).

Necropsy of *PPa1^hpt/hpt^* revealed massively enlarged spleens with ∼3-fold increase in cellularity (Fig. 2a and b). The enlarged spleens contained high frequencies of hematopoietic stem and progenitor cells (HSPCs), suggesting extramedullary hematopoiesis (EMH) and a problem with hematopoiesis within the bone marrow medullary cavity (Fig. 2c-e). The long bones from *PPa1^hpt/hpt^* mice were pale compared to *PPa1^+/+^*mice and had slightly increased total cellularity (Fig. 2f and g) while retaining approximately normal frequencies of HSPCs (Fig. 2h-j). Additionally, screening of *PPa1^hpt/hpt^*mice with DEXA scanning identified decreased bone mineral density (BMD) (Fig. 2k)^14^. MicroCT of distal femurs from *PPa1^hpt/hpt^* mice showed decreased bone volume, decreased trabeculae, and increased inter-trabecular space consistent with severe osteoporosis (Fig. 2l and m).

**Figure 2:**
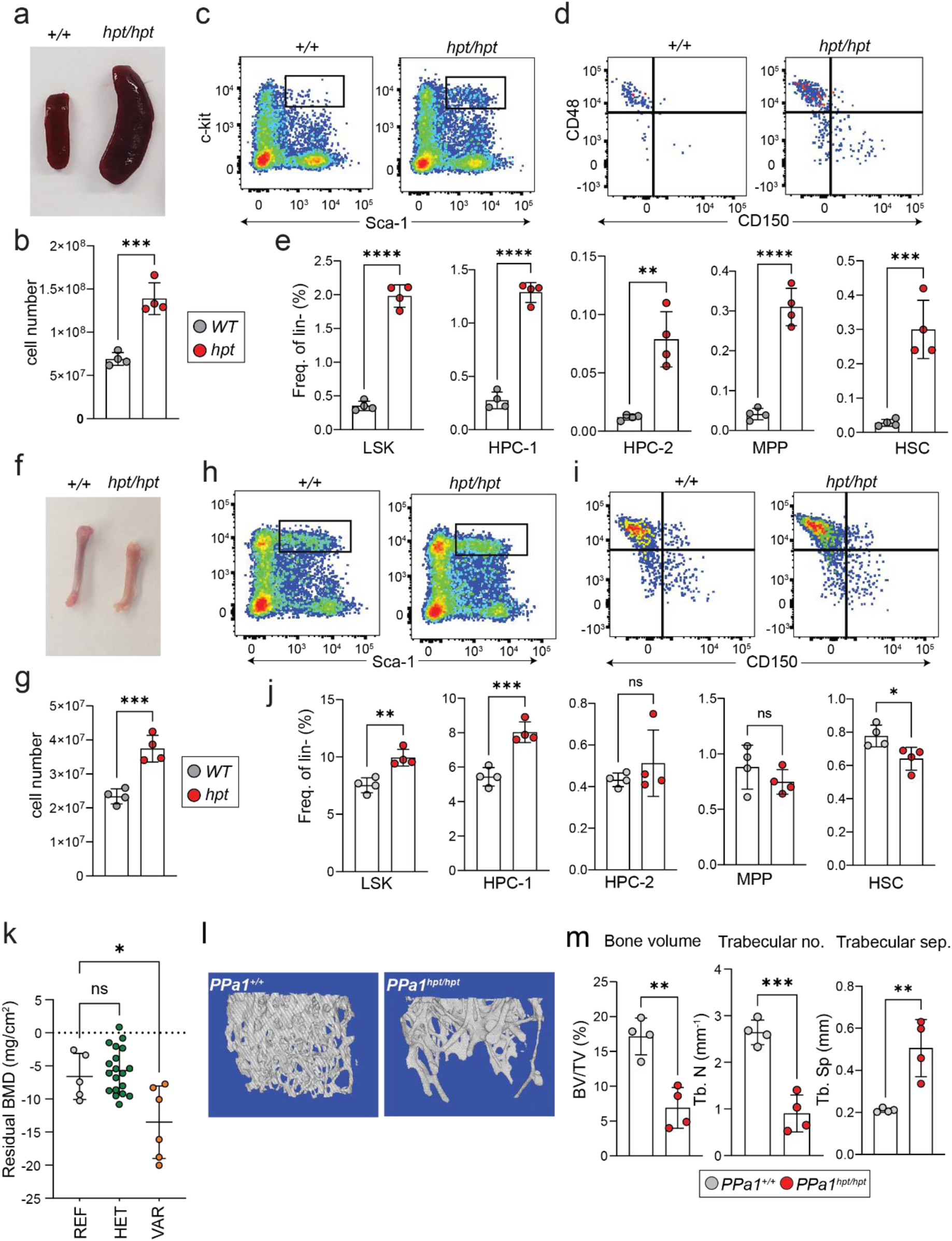
PPa1-deficiency leads to extramedullary hematopoiesis and osteoporosis. **(a)** Splenomegaly and **(b)** total spleen cell counts in *PPa1^hpt/hpt^*mice. **(c-e)** Expansion of hematopoietic progenitors in the spleens of *PPa1^hpt/hpt^* mice. **(f)** Gross bone morphology and **(g)** total bone marrow cell counts in *PPa1^hpt/hpt^* mice. **(h-j)** Frequency of hematopoietic progenitor populations in the bone marrow of *PPa1^hpt/hpt^* mice. **(k)** DEXA screening of *hotpot* pedigree shows decreased tibia bone mineral density. **(l, m)** MicroCT of distal femur of *PPa1^hpt/hpt^* mice showing osteoporosis. In all panels each symbol represents an individual mouse. Data are expressed as mean ± s.d. and significance was determined with two-tailed unpaired t-test **(b, e, g, j, m)** and one-way ANOVA with Dunnett’s multiple comparisons.

Altogether these findings indicate that *PPa1*-deficiency causes a spontaneous disease process that interferes with normal bone marrow hematopoietic differentiation and skeletal homeostasis. Interestingly, this phenotype resembled infiltrative bone marrow disorders, including leukemia, fungal infections, and lysosomal storage diseases (LSDs), in which expansion of inflammatory or transformed cell populations displaces normal hematopoietic elements and disrupts skeletal stem cell differentiation.

### Dose-dependent relationship between PPa1 expression and hematopoietic defects

Through random mutagenesis, we generated a second *PPa1* allele encoding an isoleucine-to-phenylalanine substitution at position 91 (I91F). Homozygous I91F mice (*PPa1^I91F/I91F^*) displayed normal WBC counts but exhibited anemia with anisocytosis (Fig. S2a, b). No differences in bone mineral density were detected in *PPa1^I91F/I91F^* mice (Fig. S2c).

Structural modeling indicated that the I91 residue lies outside the enzyme active site (Fig. S2d), and AlphaMissense predicted the I91F substitution to be damaging (pathogenicity score: 0.841). *PPa1^I91F/I91F^*mice showed intermediate PPa1 protein expression in bone marrow compared to WT and *PPa1^hpt/hpt^* mice (Fig. S2e).

To examine the relationship between PPa1 dosage and hematopoietic function, we generated an allelic series by intercrossing *PPa1^I91F/I91F^* mice with *PPa1^+/-^* and *PPa1^hpt/hpt^* mice (Fig. S2f). As PPa1 levels decreased, splenomegaly increased and peripheral neutropenia, anemia, and anisocytosis worsened (Fig. S2g). These findings establish a dosage-sensitive requirement for PPa1 in maintaining normal hematopoiesis.

### An atypical myeloid cell type infiltrates the bone marrow of PPa1-deficient mice

Histological analysis of *PPa1^hpt/hpt^* femurs revealed bone marrow infiltration of a unique histiocytic cell population harboring large amounts of pink, lamellar cytoplasm (Fig. 3a, b). These cells stained positively for PAS and F4/80, indicating that they contained glycosylated lipid or protein species and were of myeloid origin (Fig. 3b). These cells morphologically resembled lipid-laden macrophages that are frequently observed infiltrating tissues of patients with LSDs, such as foam cells in Niemann-Pick Disease or Gaucher cells in Gaucher disease^15^. TEM analysis of *PPa1^hpt/hpt^* femurs showed bone marrow cells containing electron dense tubular lysosomes that resembled prior models of LSD macrophage dysfunction (Fig. 3b)^16^.

**Figure 3:**
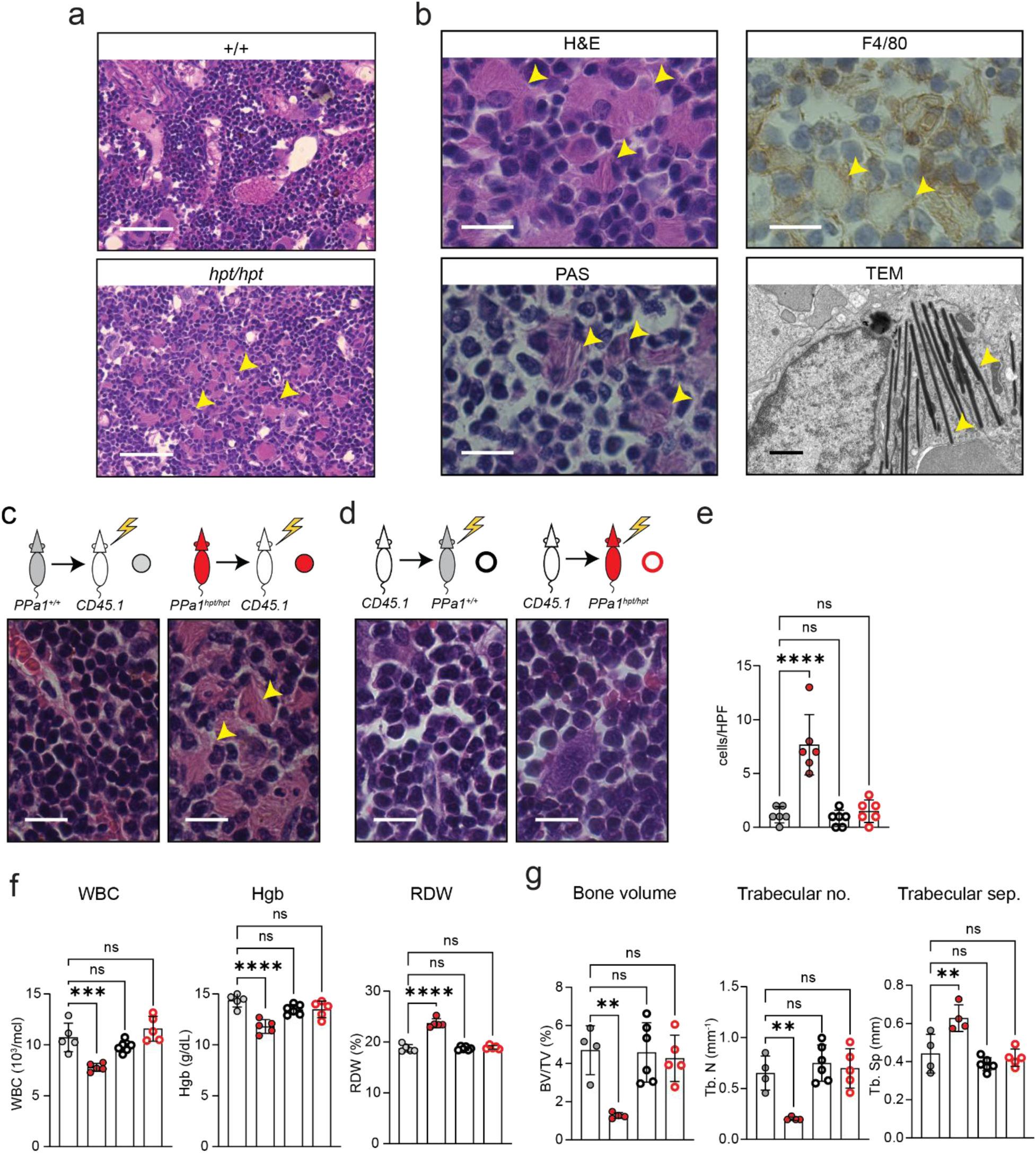
Bone marrow infiltrative disease in PPa1-deficient mice. **(a)** 40X H&E stain of bone marrow from *PPa1^+/+^* and *PPa1^hpt/hpt^* mice. Yellow arrows indicate infiltrating cells with large amounts of pink cytoplasm in *PPa1^hpt/hpt^* bone marrow. Scale = 100 µm. **(b)** 100X magnification of infiltrating cells from *PPa1^hpt/hpt^* bone marrow using the indicated stains and 8000X magnification of *PPa1^hpt/hpt^* bone marrow using TEM. Yellow arrows indicate infiltrating cells (light microscopy) and tubular lysosomes (TEM). Scale for light microscopy = 20 µm. Scale for TEM = 1 µm. **(c, d)** Reciprocal bone marrow transplantation of *PPa1^+/+^* or *PPa1^hpt/hpt^*bone marrow into lethally irradiated CD45.1 mice and vice versa with representative 100X H&E sections from recipients 40 weeks post-transplant. Scale = 20 µm. **(e)** Quantification of infiltrating macrophages across 6 random 100X fields of recipient mice. Each symbol represents the number of infiltrating macrophages in a high-power field (hpf). **(f)** WBC counts and RBC indices from the peripheral blood of transplanted mice at 12 weeks post-transplant. **(g)** MicroCT analysis of the femurs from the transplanted mice at 40 weeks post-transplant. In **(f, g)** each symbol represents an individual recipient mouse and results are representative of two independent reciprocal transplant experiments. Data are expressed as mean ± s.d. and significance was determined with one-way ANOVA with Dunnett’s multiple comparisons **(e-g)**.

### Hematologic, bone, and infiltrative phenotypes are hematopoietic intrinsic

PPi secreted from chondrocytes and osteoblasts is a physiologic inhibitor of bone mineralization where it directly antagonizes the deposition of hydroxyapatite^17, 18^. Additionally, *PPa1* is expressed in chondrocytes and osteoblasts suggesting that intracellular control of PPi levels may influence bone matrix mineralization^19–21^. Therefore, decreased PPa1 activity in these extra-hematopoietic cell populations could cause an osteoporosis phenotype. Alternatively, a hematopoietic-intrinsic bone marrow infiltrative disease could also lead to decreased mineralization as is frequently seen in LSDs^22^.

To resolve these two possibilities, we performed reciprocal bone marrow transplantation experiments (Fig. 3c, d). Reconstitution of lethally irradiated *PPa1^+/+^* mice with *PPa1^hpt/hpt^*bone marrow induced the characteristic infiltrative disease in the recipients (Fig. 3c), whereas transplantation of *PPa1^+/+^* bone marrow into *PPa1^hpt/hpt^* mice significantly reduced the infiltrative disease burden (Fig. 3d, e). Additionally, transplantation of *PPa1^hpt/hpt^* bone marrow recapitulated the hematologic defects observed in *PPa1^hpt/hpt^* mice whereas these were cured upon transplantation of *PPa1^+/+^* bone marrow into *PPa1^hpt/hpt^* mice (Fig. 3f).

MicroCT of distal femurs from *PPa1^+/+^* mice 40 weeks after receiving *PPa1^hpt/hpt^* bone marrow demonstrated a striking loss of mineralized bone (Fig. 3g). These skeletal deficiencies were normalized in *PPa1^hpt/hpt^* recipients 40 weeks after receiving *PPa1^+/+^* bone marrow (Fig. 3g). Together, these results show that both the hematopoietic and skeletal phenotypes observed in PPa1-deficient mice are transmitted by cells of hematopoietic origin.

### Identification of unique myeloid populations in *PPa1*-deficient bone marrow

To identify the molecular characteristics of the infiltrative myeloid cell population seen in *PPa1*-deficient bone marrow, we performed single-cell RNA sequencing (scRNA-seq) on bone marrow mononuclear cells pooled from three *PPa1^hpt/hpt^* mice and three *PPa1^hpt/+^*littermate controls, which do not display bone or hematologic phenotypes. After quality control and normalization, we aggregated the data from both genotypes and performed unsupervised clustering in Seurat, yielding 21 transcriptionally distinct clusters visualized by uniform manifold approximation and projection (UMAP) (Fig. 4a and b). Analysis of highly expressed genes within each cluster revealed all major hematopoietic lineages expected in bone marrow.

**Figure 4:**
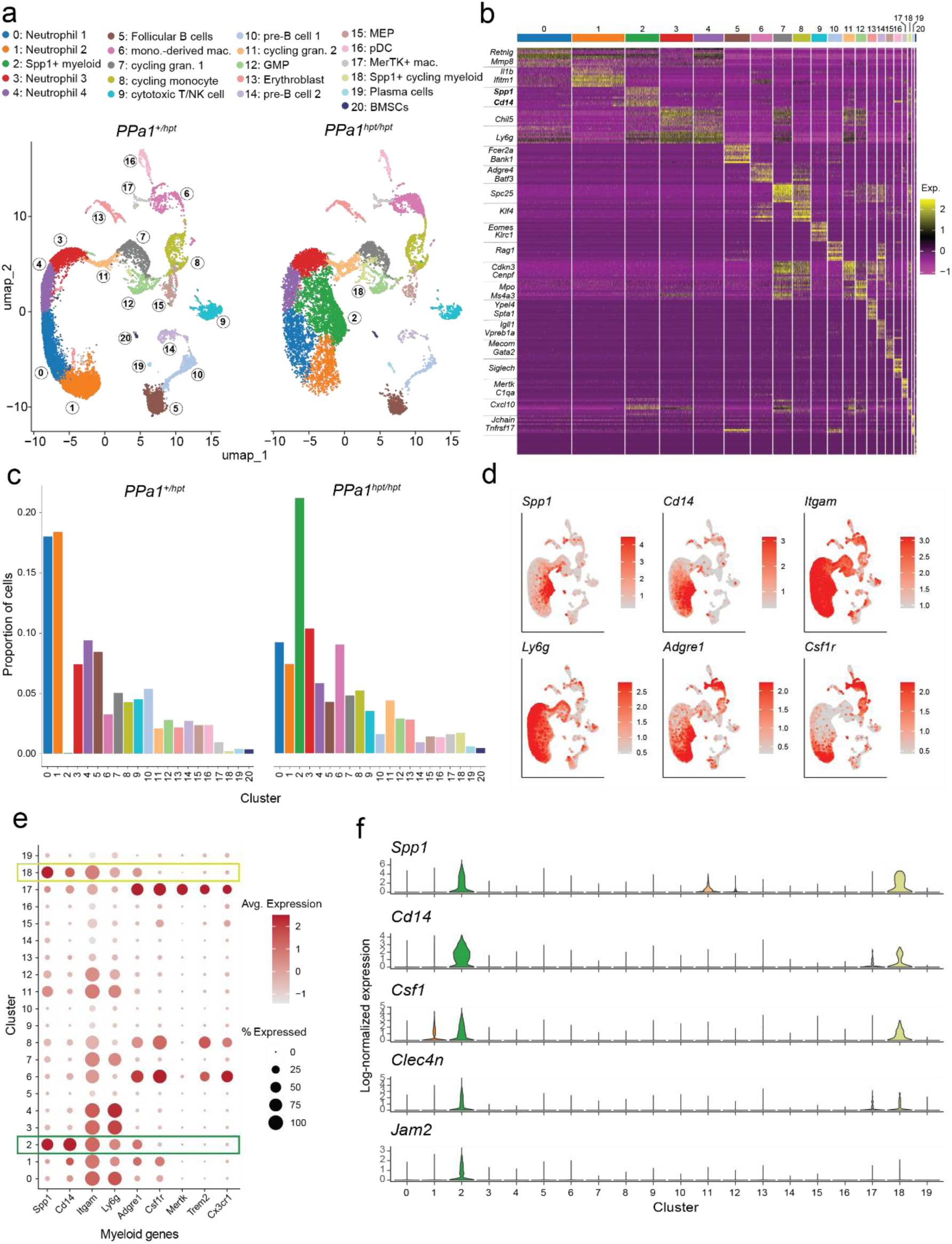
scRNA-seq identifies unique Spp1+ myeloid populations in *PPa1^hpt/hpt^*bone marrow. **(a)** UMAP projection of merged bone marrow scRNA-seq data comprising 21,703 cells from *PPa1^hpt/+^* mice and 15,802 cells from *PPa1^hpt/hpt^* mice. Each dot represents an individual cell. A total of 21 clusters (0–20), corresponding to major bone marrow lineages, were identified and color-coded. **(b)** Heatmap showing the top 10 most highly expressed genes in each cluster, with cluster-defining genes indicated. A complete list of cluster-defining genes is provided in Table S1. **(c)** Split bar graph depicting the relative frequency of each cluster by genotype. **(d)** Feature plots showing expression of selected genes projected onto the merged UMAP of bone marrow scRNA-seq data from *PPa1^hpt/+^* and *PPa1^hpt/hpt^* mice. **(e)** Dot plot displaying expression of major myeloid marker genes across clusters 0–19. **(f)** Violin plots showing expression of selected genes across clusters 0–19.

Most cells in the dataset were of myeloid origin. Clusters 0, 1, 3, and 4 corresponded to neutrophils (*Ly6g*, *Mmp8*, *Mmp9*, *Retnlg*), while clusters 7 and 11 contained cycling granulocyte precursors (*Cebpe*, *Cenpf*, *Cdkn3*). Cluster 12 represented granulocyte–monocyte progenitors (GMPs; *Elane*, *Mpo*, *Ms4a3*). Monocyte–macrophage populations included Trem2^+^ monocyte-derived macrophages (cluster 6), MerTK^+^ resident macrophages (cluster 17), and a cycling macrophage population (cluster 8). Cluster 16 corresponded to plasmacytoid dendritic cells (*Siglech*, *Tcf4*, *Irf8*) (Fig. 4b and Table S1).

Non-myeloid populations were also detected. Clusters 10 and 14 contained pre–B cells (*Vpreb3*, *Pax5*, *Il7r*), cluster 5 contained mature recirculating B cells (*Cr2*, *Fcer2a*, *Ms4a1*), and cluster 19 contained plasma cells (*Jchain*, *Tnfrsf17*). Cluster 9 contained cytotoxic T/NK cells (*Eomes*, *Klrc2*, *Cd3e*), while clusters 13 (*Ypel4*, *Spta1*, *Ermap*) and 15 (*Mecom*, *Gata2*, *Meis1*) corresponded to erythroblasts and megakaryocyte–erythroid progenitors (MEPs), respectively. A small population of bone marrow mesenchymal stromal cells (*Cxcl12*, *Kitl*, *Lepr*) was present in cluster 20.

*PPa1^hpt/hpt^* mice showed a striking expansion of cluster 2, a myeloid population expressing high levels of *Cd14* and *Spp1* (osteopontin, OPN) (Fig. 4a and c). Additionally, they also uniquely harbored a smaller cluster of cycling Spp1^+^ cells (cluster 18). The expansion of cluster 2 occurred at the expense of cells in clusters 0 and 1. *PPa1^hpt/hpt^* mice also displayed increased monocyte-derived macrophages in cluster 6 but similar proportions of MerTK^+^ resident macrophages in cluster 17. The proportions of GMPs in cluster 12 and cycling myeloid progenitors in clusters 8, 7, and 11 were comparable between *PPa1^hpt/+^* and *PPa1^hpt/hpt^*bone marrow. Consistent with the peripheral B cell deficiency in *PPa1^hpt/hpt^* mice, B cell progenitors and mature B cells were reduced. Although erythroblast frequencies were similar between genotypes, MEPs were selectively decreased in *PPa1^hpt/hpt^* bone marrow. Thus, PPa1 deficiency leads to expansion of unique Spp1^+^ myeloid populations and suppression of normal mature neutrophils, B cells, and erythroid precursors.

### CD14^+^Spp1^+^ myleoid cells have features of inflammatory neutrophils and monocytes

The CD14^+^Spp1^+^ cluster unique to *PPa1*^hpt/hpt^ mice represented a transcriptionally distinct myeloid population. It expressed *Itgam* (CD11b), confirming myeloid identity, and showed intermediate expression of the macrophage marker *Adgre1* (F4/80) and the neutrophil marker *Ly6g* (Fig. 4d). Notably, cluster 2 lacked expression of canonical macrophage markers like *Csf1r*, *Mertk*, *Trem2*, and *Cx3cr1*, but showed high expression of the monocyte-specific gene *Cd14* (Fig. 4d and e). Additional markers that distinguished cluster 2 from other bone marrow myeloid cells included *Csf1, Clec4n* (Dectin-2), and *Jam2* (Fig. 4f). Together, these features show that the CD14^+^Spp1^+^ cluster represents an atypical inflammatory myeloid population with markers of both monocytes and granulocytes.

To further define the CD14⁺Spp1⁺ myeloid population, we analyzed inflammatory gene expression signatures in the scRNA-seq data (Fig. 5a). In addition to *Spp1*, myeloid cells in clusters 2 and 18 expressed high levels of the inflammatory cytokines *Csf1*, *Ccl3*, and *Ccl4*. These populations did not exhibit a type I interferon gene signature. While Il1b expression was detectable, there was no evidence of a robust inflammasome-associated transcriptional program. Additionally, although these cells expressed intermediate levels of Tnf, they did not conform to classical M1 or M2 polarization states.

**Figure 5:**
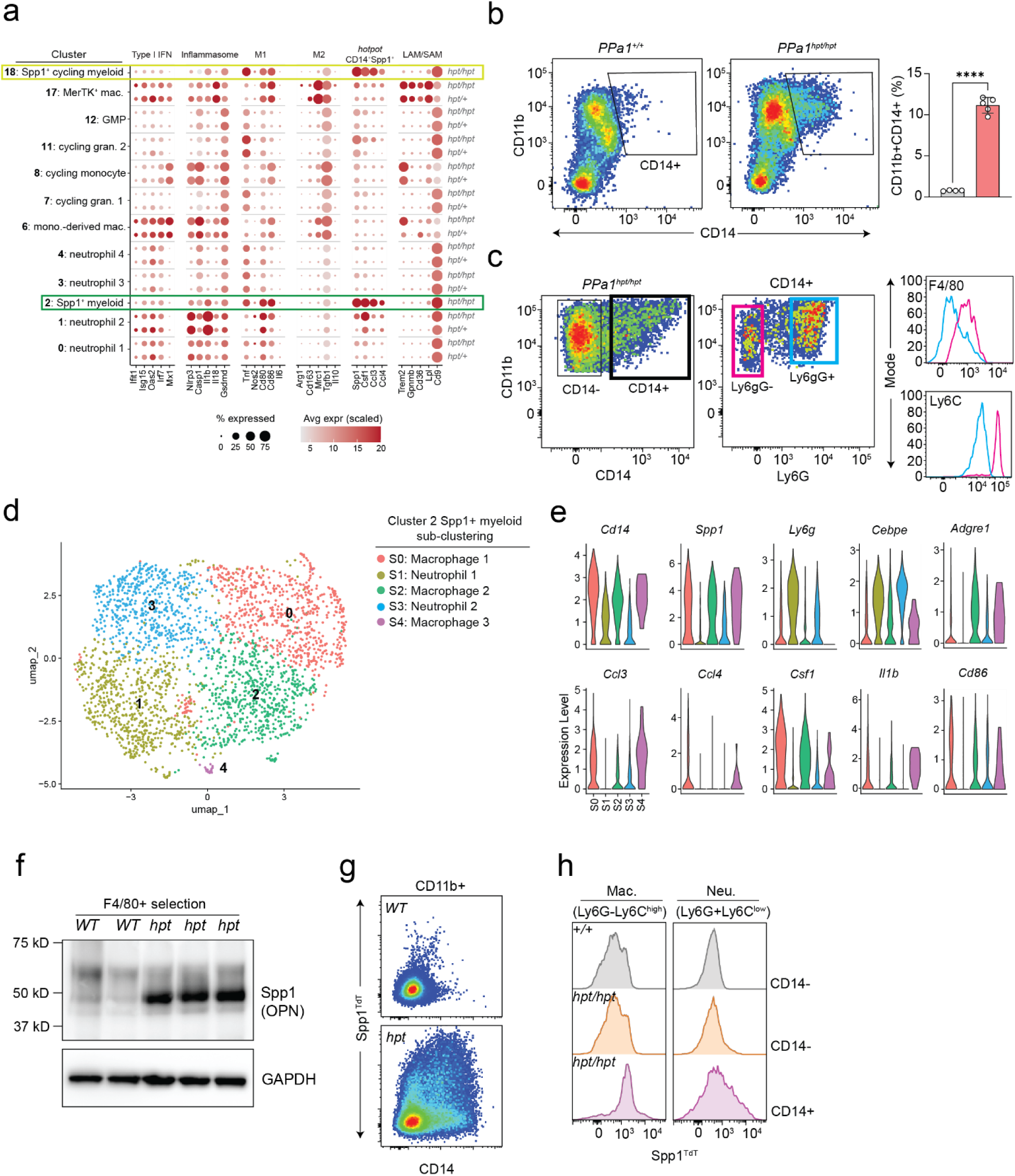
Validation and characterization of CD14⁺Spp1⁺ myeloid cells in *PPa1^hpt/hpt^*bone marrow. **(a)** Dot plot showing expression of inflammatory genes across myeloid clusters, split by genotype. Clusters 2 and 18 were selectively enriched in *PPa1^hpt/hpt^* mice. **(b)** Flow cytometric quantification showing expansion of the CD11b⁺CD14⁺ myeloid population in *PPa1^hpt/hpt^* bone marrow. Each symbol represents an individual mouse. **(c)** The CD11b⁺CD14⁺ population comprises two subsets: a Ly6G⁺F4/80⁻Ly6C^low^ neutrophil-like population and a Ly6G⁻F4/80⁺Ly6C^high^ macrophage-like population. **(d)** UMAP of reclustered CD14⁺Spp1⁺ myeloid cells (original cluster 2). Each dot represents a single cell. **(e)** Violin plots showing expression of selected genes across subclusters of Spp1⁺ myeloid cells. **(f)** Immunoblot analysis of Spp1 protein expression in F4/80⁺ bone marrow macrophages. **(g, h)** Spp1 reporter expression across bone marrow myeloid cell subsets. Data are expressed as mean ± s.d. and significance was determined with two-tailed unpaired t-test **(b)**.

Notably, we did not observe strong enrichment of inflammatory gene signatures in conventional *PPa1^hpt/hpt^* macrophage and neutrophil populations, demonstrating that PPa1-deficiency does not broadly skew bone marrow myeloid cells toward an inflammatory phenotype (Fig. 5a).

Spp1⁺ macrophages have been described in multiple metabolic and fibrosing disease states where they are called lipid-associated macrophages (LAMs) or scar-associated macrophages (SAMs)^23–25^. These cells are typically characterized by co-expression of *Trem2*, *Gpnmb*, *Cd36*, *Lpl*, and *Cd9*. We did not detect specific expression of these transcripts within Spp1^+^ cell populations in our dataset, indicating that PPa1-deficient CD14⁺Spp1⁺ myeloid cells are transcriptionally distinct from reported LAM/SAM macrophage populations (Fig. 5a).

### Phenotypic validation of CD14⁺Spp1⁺ myeloid cells in PPa1-deficient bone marrow

We next sought to validate the presence of CD14⁺Spp1⁺ myeloid cells within the bone marrow. We first assessed CD14 expression within bone marrow myeloid subsets (Fig. 5b). *PPa1^hpt/hpt^* mice contained a prominent CD11b⁺CD14^hi^ population that was absent in PPa1^+/+^ bone marrow. This population contained both Ly6G⁺ and Ly6G^-^ subsets, suggesting contributions from granulocyte-related and monocyte-derived cells (Fig. 5c). The Ly6G^-^ subset expressed high Ly6C and intermediate F4/80, further supporting monocyte/macrophage origins.

To further resolve the heterogeneity within CD14^+^Spp1+ myeloid cells, we reclustered this population following normalization, data scaling, and regression of mitochondrial, ribosomal, and cell cycle-associated genes. This analysis identified five transcriptionally distinct subclusters on UMAP (Fig. 5d). Although all subclusters expressed *Cd14*, substantial heterogeneity was evident, including discrete neutrophil-like and macrophage-like populations (Table S2). Subclusters 1 and 3 exhibited a granulocytic identity, marked by high expression of *Ly6g* and *Cebpe*, and low-to-moderate *Spp1* expression (Fig. 5e). In contrast, subclusters 0, 2, and 4 displayed more prominent macrophage features. Subcluster 0, particularly, was enriched for an inflammatory Spp1⁺ program with elevated expression of Ccl3, Ccl4, Il1b, and Csf1.

To confirm Spp1 expression within PPa1-deficient macrophages, we performed bead-based sorting of F4/80⁺ cells from *PPa1^hpt/hpt^* and *PPa1^+/+^* bone marrow followed by immunoblotting (Fig. 5f). *PPa1^hpt/hpt^*macrophages showed markedly elevated Spp1 protein, whereas *PPa1^+/+^*cells showed little to no expression. We next crossed *PPa1^hpt/hpt^*mice to an Spp1-IRES-tdTomato (Spp1^TdT^) reporter strain to detect Spp1⁺ bone marrow myeloid cells^25^. This revealed a unique population of Spp1^TdT^⁺;CD14⁺ myeloid cells in *PPa1^hpt/hpt^* bone marrow (Fig. 5g). Spp1-TdT expression was highest in *PPa1^hpt/hpt^* CD14^+^Ly6G^-^Ly6C^high^ macrophages compared to *PPa1^hpt/hpt^* or *PPa1^+/+^* CD14^-^Ly6G^-^Ly6C^high^ macrophages (Fig. 5h). Minimal Spp1^TdT^ reporter was seen in Ly6G^+^ neutrophil-like populations. Together, these findings establish that PPa1 insufficiency drives spontaneous expansion of CD14⁺Spp1⁺ inflammatory macrophages within the bone marrow.

### PPa1-deficiency causes bone marrow sphingolipidosis and myeloid lysosomal dysfunction

The infiltrating macrophages in *PPa1^hpt/hpt^* mice morphologically resembled lipid-laden macrophages that accumulate in the tissues of patients with LSDs. In particular, the pink lamellar cytoplasm of infiltrating bone marrow histiocytes (“crinkled paper cytoplasm”) in *PPa1^hpt/hpt^* mice resembled Gaucher cells, a distinct population of macrophages found in patients with Gaucher Disease (GD) who carry loss-of-function mutations in *glucocerebrosidase* (*GBA*)^26^. Gaucher cells accumulate undigested glycosphingolipids within their lysosomes and their infiltration into bone marrow is linked to defective hematopoiesis and skeletal homeostasis, although the precise mechanism for how these occur is not clear^27^. Additionally, high levels of circulating Spp1 are characteristic of GD and its level inversely corresponds to enzyme replacement efficacy^28^.

Based on the phenotypic similarity to GD, we speculated that PPa1-deficient mice would exhibit sphingolipid accumulation within their bone marrow. *PPa1^hpt/hpt^* mice displayed significantly elevated levels of long chain glucosyl ceramides compared to *PPa1^+/+^* mice (Fig. 6a and b). Additionally, we noted a marked increase in sphingosine levels in *PPa1^hpt/hpt^*mice without a corresponding rise in its downstream metabolite, sphingosine-1-phosphate (S1P) (Fig. 6c). This finding parallels observations in GBA1 knockout mice, where glucosyl-sphingosine accumulates within lysosomes, subsequently diffuses into the cytoplasm, and is catabolized into sphingosine by cytoplasmic GBA2^29^. However, despite sharing several characteristics with GBA1-deficient mice and humans, PPa1-deficient macrophages showed increased levels of GBA1 protein expression and similar levels of enzyme activity compared to WT cells (Fig. 6d).

**Figure 6:**
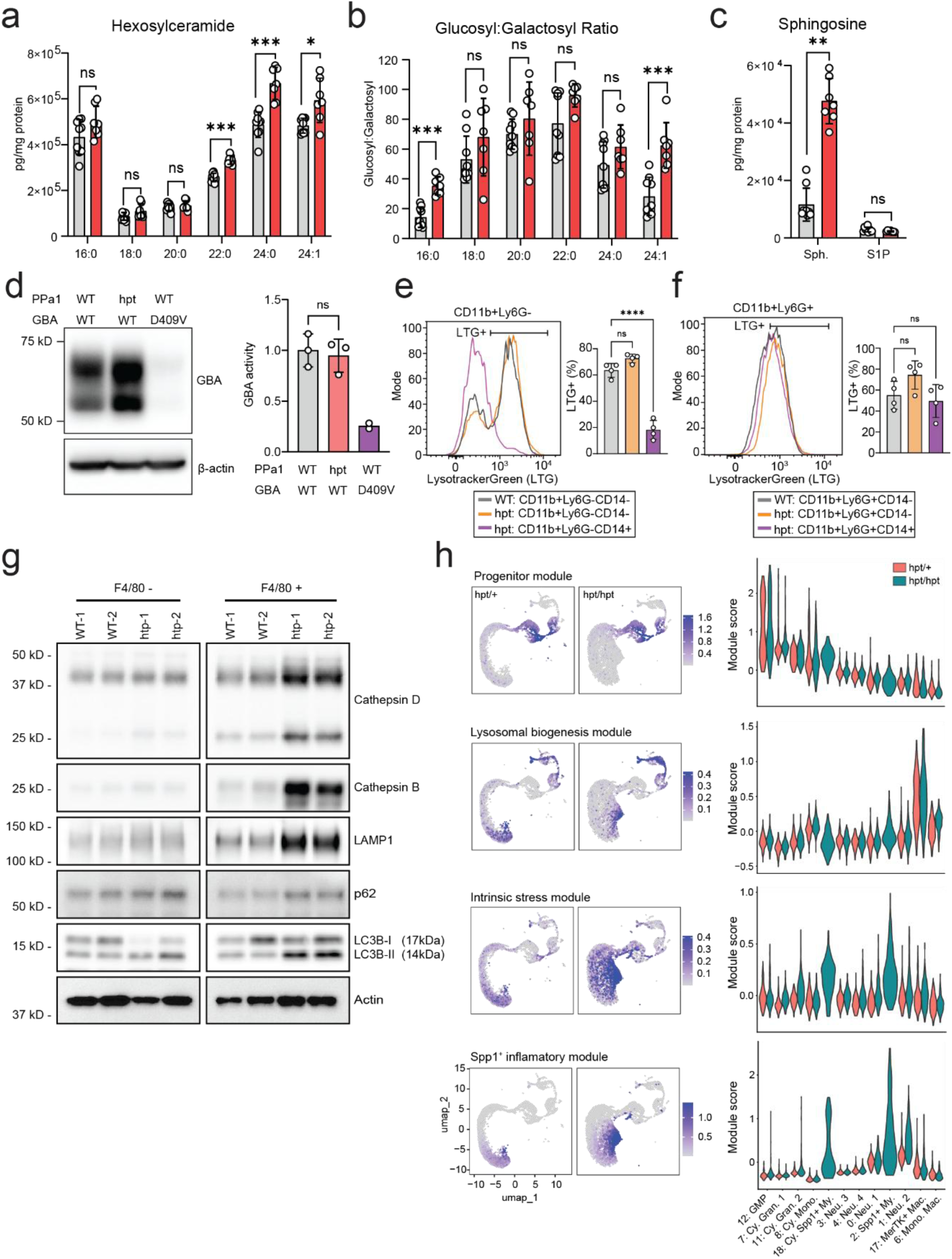
PPa1 deficiency induces bone marrow sphingolipidosis and macrophage lysosomal dysfunction. **(a–c)** Quantitative sphingolipid analysis of bulk bone marrow from *PPa1^+/+^* (gray) and *PPa1^hpt/hpt^*(red) mice. Each symbol represents an individual mouse. **(d)** Immunoblot analysis of GBA expression in F4/80⁺ macrophages from *PPa1^hpt/hpt^*and *GBA^D409V^* mice and GBA enzymatic activity in F4/80⁺ macrophages from the indicated genotypes, normalized to *PPa1^+/+^* controls. Each symbol represents the mean of three technical replicates from bone marrow of an individual mouse. **(e, f)** LysoTracker Green (LTG) staining in bone marrow myeloid subsets from *PPa1^+/+^* and *PPa1^hpt/hpt^* mice. Each symbol represents an individual mouse. **(g)** Immunoblot analysis of lysosomal and autophagy markers in F4/80⁺ bone marrow macrophages. **(h)** Feature and violin plots showing expression of the indicated gene modules across myeloid populations, split by genotype. Myeloid subsets are ordered from least to most mature based on progenitor score. Module scores were calculated using curated gene sets (Table S3). Data are expressed as mean ± s.d. and significance was determined with Mann-Whitney test **(a-c)**, two-tailed unpaired t-test **(d)**, and one-way ANOVA with Tukey’s multiple comparisons **(e, f).**

To further characterize lysosomal defects in PPa1-deficient myeloid populations, we stained *PPa1^hpt/hpt^* and *PPa1^+/+^*bone marrow cells with Lysotracker Green (LTG) to label acidic organelles (Fig. 6e and f). CD14^+^Ly6G^-^Ly6C^high^ cells from *PPa1^hpt/hpt^*mice showed strikingly reduced LTG fluorescence compared to CD14^-^Ly6G^-^Ly6C^high^ cells from either *PPa1^hpt/hpt^* or *PPa1^+/+^* mice. Additionally, there was no difference in LTG intensity between Ly6G^+^ neutrophil-like populations from either *PPa1^hpt/hpt^* or *PPa1^+/+^* mice. Together these data show that PPa1-deficient CD14^+^Spp1^+^ macrophage populations have a specific defect in lysosomal acidification that is not seen in WT monocyte/macrophage populations or in PPa1-deficient granulocyte populations.

### PPa1 does not localize to the lysosome under steady state conditions in bulk bone marrow

High concentrations of PPi can buffer protons and chelate divalent cations. We hypothesized that PPa1 might act as a lysosomal hydrolase that degrades PPi ingested by phagocytes to facilitate normal lysosomal function. We isolated lysosomes from the bone marrow of transgenic mice expressing an HA-tagged lysosomal resident protein, TMEM192, driven by Vav-iCre from the Rosa26 locus in hematopoietic cells (Lyso-IP mice) (Fig. S3a). PPa1 did not co-fractionate with the autophagosome/lysosomal markers Lamp1, LC3b-II, and Cathepsin B. While these data do not support a primary role for it as a lysosomal hydrolase, they cannot rule out PPa1’s association with the lysosomal machinery in rare subsets of myeloid cells.

### PPa1 deficiency induces metabolic stress in mature myeloid cells without compensatory lysosomal biogenesis

In sorted F4/80⁺ macrophages from *PPa1^hpt/hpt^* mice, we observed enrichment of several lysosomal and autophagy-associated proteins, including LAMP1, CtsB, CtsD, LC3B-II, and p62 (Fig. 6g). Notably, these phenotypes were restricted to cells isolated directly from the bone marrow. We did not detect major changes in Spp1 expression or lysosomal proteins in bone marrow-derived macrophages (BMDMs) differentiated in vitro under CSF1 stimulation (Fig. S3b).

We sought to determine whether the accumulation of lysosomal proteins with PPa1-deficient bone marrow macrophages reflected increased lysosomal biogenesis or, alternatively, impaired lysosomal maturation or resolution. In parallel, we aimed to define whether alterations in lysosomal pathways, intrinsic cellular stress, and inflammatory programming driven by PPa1 deficiency arise early during myeloid development or emerge in mature myeloid populations.

To address this, we ordered bone marrow myeloid clusters by differentiation state using a composite progenitor score integrating GMP and cell-cycle gene signatures (Fig. 6h, Table S3). As expected, cluster 12 (GMP) exhibited the highest progenitor score, followed by proliferating granulocyte and monocyte precursors (clusters 7, 11, and 8), the cycling CD14⁺Spp1⁺ myeloid population (cluster 18), and then terminally differentiated populations including mature neutrophils (clusters 0, 1, 3, and 4), monocytes (clusters 6 and 17), and CD14⁺Spp1⁺ myeloid cells (cluster 2).

The PPa1-deficiency-associated inflammatory signature was not strongly induced in progenitor populations or in canonical mature monocytes, macrophages, or most neutrophil subsets (Fig. 6h, Table S3). Instead, this signature was selectively enriched in Spp1^+^ myeloid populations (cluster 18 and cluster 2), with weaker induction in neutrophils (cluster 1). Notably, CD14⁺Spp1⁺ myeloid cells in PPa1-deficient mice did not exhibit robust activation of lysosomal biogenesis programs, arguing against compensatory lysosome expansion. However, we observed strong and selective induction of intrinsic cellular stress response programs in both Spp1^+^ myeloid populations (clusters 18 and 2).

Taken together, these transcriptional and proteomic data indicate that PPa1 deficiency leads to metabolic stress in specific myeloid subpopulations where lysosomal maturation is compromised and a distinct pro-inflammatory state emerges.

## DISSCUSION

Here, we present the first mouse model of systemic PPa1 deficiency. Although PPa1 has been proposed to play a housekeeping role that is necessary for cell growth, proliferation, and metabolism, we observed a surprisingly specific syndrome that recapitulates key features of LSDs. PPa1-deficient mice exhibited defective medullary hematopoiesis, extensive extramedullary hematopoiesis, accumulation of glycosphingolipids in the bone marrow, and impaired bone mineralization. These abnormalities were associated with marked infiltration of lipid-laden macrophages into the bone marrow. Further characterization revealed bone marrow infiltration of macrophages displaying a distinct inflammatory program marked by Spp1 and CD14 expression, along with evidence of lysosomal dysfunction and metabolic stress. Together, these findings uncover a previously unrecognized role for PPa1 in promoting lysosomal homeostasis within the myeloid compartment.

A central question arising from this work is how PPa1 regulates lysosomal function in myeloid cells. We found that PPa1 is predominantly cytosolic and does not co-fractionate with lysosomes, arguing against a direct role as a lysosomal hydrolase. Given the central role of pyrophosphate (PPi) hydrolysis in multiple metabolic pathways, one possibility is that loss of PPa1 activity induces cellular metabolic stress that secondarily impairs lysosomal function. Consistent with this idea, loss-of-function mutations in the yeast PPi hydrolase IPP1 lead to NAD depletion and profound cellular stress, accompanied by marked vacuolization^5^. This phenotype may be analogous to the lysosomal abnormalities observed in PPa1-deficient macrophages. In further support of increased cellular stress during PPa1 deficiency, we observed enrichment of intrinsic stress-response gene programs in Spp1⁺ myeloid cells. Whether this stress response is causal for lysosomal dysfunction or instead arises because of impaired lysosomal homeostasis remains to be determined.

Another possibility is that PPa1 has important functions outside of PPi hydrolysis. Recent studies have demonstrated that loss of PPa1 causes defective adipocyte differentiation and impaired adipocyte browning^8, 30^. Notably, these defects could be rescued by catalytically inactive but structurally stable PPa1. Therefore, it is possible that key PPa1 functions may depend on protein stability rather than enzymatic activity and would suggest a scaffolding role that supports other effector proteins. This model may account for why homozygous human PPa1 mutations that abolish enzyme activity, but preserve significant protein levels, are not associated with severe clinical phenotypes^9^. Future experiments that measure the effects of catalytically inactive yet stable PPa1 variants on myeloid differentiation, along with identification of PPa1-interacting partners, will be critical for defining the molecular basis of PPa1 function.

Transcriptomic analysis of PPa1-deficient bone marrow identified a distinct population of inflammatory CD14⁺Spp1⁺ myeloid cells that lacked expression of several canonical macrophage markers. We found that the CD14^+^ myeloid population contained two phenotypically distinct subsets: a granulocyte cell type that expressed Ly6G and low levels of Ly6C, and a macrophage cell type that expressed F4/80 and high levels of Ly6C. It was the CD14^+^ macrophage subset that exhibited high Spp1 expression, markedly reduced LysoTracker staining, and increased transcriptional expression of cellular stress genes. Notably, in vitro differentiation of BMDMs with CSF1 failed to recapitulate the molecular features observed in endogenous CD14⁺ macrophages. Moreover, lysosomal defects and heightened inflammatory gene programs were not observed in other PPa1-deficient macrophage populations, including CD14⁻ monocyte-derived macrophages and MerTK⁺ macrophages. Collectively, these findings indicate that lysosomal dysfunction and spontaneous inflammatory programming are not universal features of PPa1-deficient macrophages and instead suggest that specific cues within the bone marrow microenvironment promote the differentiation of CD14⁺Spp1⁺ macrophages.

An important unresolved question is the developmental origin of CD14⁺Spp1⁺ macrophages in the setting of PPa1 deficiency. Single-cell RNA-sequencing did not reveal evidence of inflammatory or stress-associated programming in early myeloid progenitors, including granulocyte–monocyte progenitors (GMPs). However, the transcriptomic data identified a small population of actively cycling Spp1⁺ myeloid cells (cluster 18) that exhibited a higher progenitor score than the larger, more mature CD14⁺Spp1⁺ population (cluster 2). One possibility is that the developmental trajectory of Spp1⁺ macrophages is specified earlier in myeloid development, giving rise to a distinct branch of macrophage differentiation. This occurrence would contrast with current models of lysosomal storage disorders, in which inflammatory macrophage programming is considered to arise primarily from substrate accumulation in terminally differentiated macrophages. Further studies will be required to determine whether the cycling Spp1⁺ population functions as a progenitor for mature Spp1⁺ macrophages and whether this fate is driven by metabolic dysfunction or instead reflects a direct effect of PPa1 deficiency on signaling networks during myeloid development.

We observed a striking loss of bone mineralization in PPa1-deficient mice. Bone marrow transplantation experiments demonstrated that this phenotype is hematopoietic intrinsic, implicating hematopoietic-derived signals in driving bone loss. Notably, CD14⁺Spp1⁺ myeloid cells expressed high levels of Ccl3 and Csf1. As both of these cytokines are potent stimulators of osteoclast differentiation and activity, it is possible that enhanced bone resorption underlies the reduced bone density in PPa1-deficient mice. Alternatively, infiltrative myeloid populations may remodel the bone marrow niche in ways that suppress osteogenic stromal populations, similar to what is described in malignant bone marrow infiltrative disorders^19^. Given that skeletal abnormalities are a hallmark of several LSDs^22^, transcriptional and functional analysis of stromal populations in PPa1-deficient bone marrow will be essential to understanding how myeloid-driven inflammation and lysosomal dysfunction converge to disrupt bone homeostasis.

Overall, our findings identify PPa1 as a previously unrecognized regulator of lysosomal and metabolic homeostasis in myeloid cells and present a highly penetrant model to study the effects of LSDs on the hematopoietic and skeletal systems.

## METHODS

### Mouse strains

Mice were housed under specific pathogen–free conditions and fed a standard chow diet at the University of Texas Southwestern Medical Center. All animal experiments were conducted in accordance with institutionally approved protocols. ENU mutagenesis, strategic breeding of mutagenized mice, phenotypic screening, and automated meiotic mapping were performed as previously described^10^. C57BL/6 CD45.1, Spp1^TdT^, and GBA^D409V^ mice were obtained from The Jackson Laboratory. Both male and female mice aged 8–16 weeks were used for most experiments. For reciprocal BMT experiments, female mice were used as donors and recipients. Recipient mice were allowed to reconstitute for 40 weeks to allow stabilization of bone phenotypes prior to skeletal assessments.

### Sphingolipid extraction, LC-MS/MS sphingolipid quantification, and determination of hexosyl-ceramide ratios

Bone marrow samples were lysed in 500 µL ice-cold PBS using a probe sonicator (30% power; 10 cycles of 5 s on/5 s off) while maintained on ice. Fifty microliters of lysate was reserved for protein quantification by BCA assay (Pierce). The remaining lysate (400 µL) was quenched with 2.0 mL organic extraction solvent (isopropanol:ethyl acetate, 15:85 v/v) in borosilicate glass tubes. Internal standards (20 µL; Avanti Polar Lipids) were added immediately, followed by 1.5 mL HPLC-grade water. Two-phase extraction was performed, and organic phases were collected, combined, and dried under nitrogen at 40 °C. Dried residues were reconstituted in 200 µL methanol for ceramide and sphingoid base analysis. Samples were analyzed using a Shimadzu LCMS-8060 triple quadrupole mass spectrometer coupled to a Nexera X2 UHPLC system. Ceramides and sphingoid bases were analyzed with 3 µL injections; sphingomyelins required 1 µL injections. Lipid separation was achieved using a C8 reverse-phase column under gradient elution. Metabolite concentrations were quantified using calibration curves generated from serial dilutions of authentic standards and normalized to internal standards. Following sphingolipid quantification, samples were dried and reconstituted for supercritical fluid chromatography to resolve diastereomeric hexosyl-ceramides using a Nexera UC system coupled to LCMS-8060. Diastereomeric ratios were calculated based on chromatographic peak areas.

### Protein immunoblotting

Primary cell pellets were lysed in buffer containing 1% SDS, HALT protease inhibitor cocktail, (Thermo) and benzonase. For samples analyzing GBA abundance, cells were lysed in buffer containing 1% NP-40. Protein concentrations were determined by BCA assay, and 10 µg of protein per sample was analyzed by SDS-PAGE.

### Flow cytometry

Bone marrow was flushed from femurs and tibias and samples were stained with Fc-block (Tonbo) followed by staining with the indicated fluorescent antibodies for 30 minutes on ice followed by washing in PBS. For Lysotracker Green staining, total bone marrow was equilibrated at 37 degrees for 15 minutes followed by staining with 50 nM Lysotracker for 20 minutes at 37 degrees. After incubation, cells were washed in PBS then stained with fluorescent antibodies for 30 minutes on ice. Flow cytometry data were acquired on an LSR Fortessa (BD Biosciences) and analyzed using FlowJo software.

### Lyso-IP of bulk bone marrow

Lyso-IP from bulk bone marrow was performed based on prior protocols^31, 32^. Bone marrow cells were flushed with PBS and centrifuged at 500 × g for 5 min at 4 °C. Pellets were resuspended in 1 mL ice-cold KPBS (136 mM KCl, 10 mM KH₂PO₄, pH 7.25) and centrifuged at 1,000 × g for 2 min. Cells were resuspended again in KPBS, and 100 µL was reserved as whole-cell input. The remaining cells were homogenized with 20 gentle strokes in a 2 mL homogenizer and centrifuged at 1,000 × g for 2 min. The supernatant was incubated with pre-washed anti-HA magnetic beads (Pierce) for 15 min at 4 °C with gentle rotation. Beads were washed three times with KPBS using a DynaMag-2 magnet (Invitrogen), resuspended in 2× SDS sample buffer, and boiled for 5 min. Supernatants were collected after magnetic separation for downstream analysis.

### Skeletal assessments

Distal femurs from sex-matched littermates were analyzed by microcomputed tomography (microCT) using a SkyScan 1072 X-ray Microtomograph (SkyScan, Aartselaar, Belgium) operated at 50 kV and 200 µA. A scout view of each bone was obtained to adjust the sample height and ensure that the region of interest was within the field of view. Images were acquired at an 8 µm voxel resolution with a rotation step of 0.4° between projections. Projection images were reconstructed into cross-sectional slices using NRecon software (version 1.7.4.6; SkyScan). Trabecular bone parameters were quantified using CTAn software (SkyScan). Three-dimensional renderings were generated using CTVox software (SkyScan). Trabecular measurements were performed on 200 consecutive slices of trabecular bone beginning just proximal to the distal femur growth plate using a threshold range of 79–255. Trabecular parameters were reported according to American Society for Bone and Mineral Research (ASBMR) nomenclature.

Dual energy X-ray absorptiometry (DEXA) was performed on live mice using the Faxtron UltraFocusDXA (Hologic Inc.). Mice were anesthetized with inhaled isoflurane during aquisition. Region-of-interest analysis quantified femur bone density averaged from both femurs. For Fig. 2k, animals of different age and sex were scanned, and bone density was adjusted for physiologic variation in age and sex as previously described^14^. After adjustment, residual bone density measures were used for testing statistically significant differences. For Fig. S2c, sex-matched littermates were analyzed.

### Single cell RNA sequencing

Single-cell suspensions were prepared from bone marrow of three male *PPa1^hpt/+^* and three male *PPa1^hpt/hpt^*mice. Following RBC lysis, bone marrow cells from three mice per genotype were pooled prior to library preparation. No specific enrichment for viable cells was performed; however, cell viability exceeded 90% prior to 10x capture. Samples were processed under identical conditions using standard 10x Genomics 3′ Gene Expression workflows by the UT Southwestern Microbiome Core.

Raw filtered feature-barcode matrices were generated from FASTQ files using Cell Ranger (v8.0.1; 10x Genomics) with default parameters for 3′ gene expression and aligned to the mouse reference genome mm10/GRCm38. Matrices were imported into R (v4.3.3) and analyzed using Seurat (v5.3.0). Cells were retained if they contained 500–7,000 detected features, 1,000–45,000 unique molecular identifiers (UMIs), and <5% mitochondrial gene content. After quality control filtering, 19,249 *PPa1^hpt/+^* cells and 13,835 *PPa1^hpt/hpt^* cells were retained for downstream analysis.

Counts were log-normalized (scale factor 10,000), and highly variable genes were identified using the variance-stabilizing transformation (“vst”) method. Data were scaled prior to principal component analysis. Doublets were identified independently within each genotype using DoubletFinder and removed (1,925 *PPa1^hpt/+^* and 1,384 *PPa1^hpt/hpt^* predicted doublets).

To preserve genotype-associated transcriptional differences, datasets were merged for downstream analysis rather than subjected to batch-corrected integration. Clustering was performed using a shared nearest-neighbor graph and the Louvain algorithm (resolution 0.5), and UMAP was used for visualization. Cluster marker genes were identified using Seurat’s FindAllMarkers function. Module scores were calculated using curated gene sets (Table S3).

Sequencing yielded 21,703 estimated cells with a mean of 31,017 reads per cell for the *PPa1^hpt/+^*sample and 15,802 estimated cells with a mean of 37,761 reads per cell for the *PPa1^hpt/hpt^* sample.

### Study design and statistical analysis

No statistical methods were used to pre-determine sample size. Investigators were not blinded to experimental groups during data collection. Mice were randomly allocated to experimental groups provided they met the predefined genotype, age, and sex criteria of the study.

Normality of data distribution was assessed using the Shapiro–Wilk test. For normally distributed data, differences between two groups were analyzed using paired or unpaired two-tailed Student’s *t* tests, as appropriate. For non-normally distributed data, appropriate nonparametric tests were used as indicated in the figure legends. Comparisons involving more than two groups were analyzed by analysis of variance (ANOVA) with post hoc testing as specified in the figure legends.

All statistical analyses were performed using GraphPad Prism software. Differences were considered statistically significant at *P* < 0.05. Exact *P* values and statistical tests used are provided in the corresponding figure legends.

## DATA AVAILABILITY

Raw and processed single cell RNA sequencing files analyzed in this study will be made available at the Gene Expression Omnibus (GEO) upon publication.

## ACKNOWLEDGEMENTS

This work was supported by NIH R01-AI167920 and the UT Southwestern Disease Oriented Clinical Scholars Award (to E.N.-G.). Sphingolipid analysis was performed by the UT Southwestern Metabolomics Core, which is supported by the UT Southwestern Nutrition Obesity Research Center (NORC) under NIDDK/NIH award number P30DK127984. Flow cytometry experiments were performed with assistance from the Moody Flow Cytometry Core at the Children’s Research Institute. Single cell RNA sequencing was completed by the UT Southwestern Microbiome Core and analysis of the data was aided by computational resources from the BioHPC supercomputing facility located in the Lyda Hill Department of Bioinformatics at UT Southwestern.

## AUTHOR CONTRIBUTIONS

Conceptualization (M.G, E.N.-G.); Data Curation (M.G., L.C., E.N.-G.); Formal Analysis (M.G., L.C., J.S., A.K., J.J.R., E.N.-G.); Funding Acquisition (E.N.-G.); Investigation (M.G., L.C., J.R., N.P., S.C., M.T., A.L., J.W., A.K., J.J.R., J.S., E.N.-G.); Methodology (M.G., L.C., J.R., N.P., A.K., J.J.R., E.N.-G.); Project Administration (E.N.-G.); Resources (B.B., E.N.-G.); Supervision (E.N.- G.); Validation (M.G., L.C., E.N.-G.); Visualization (M.G., L.C., E.N.-G.); Writing (original draft) (E.N.-G.).

---

**Figure S1:**
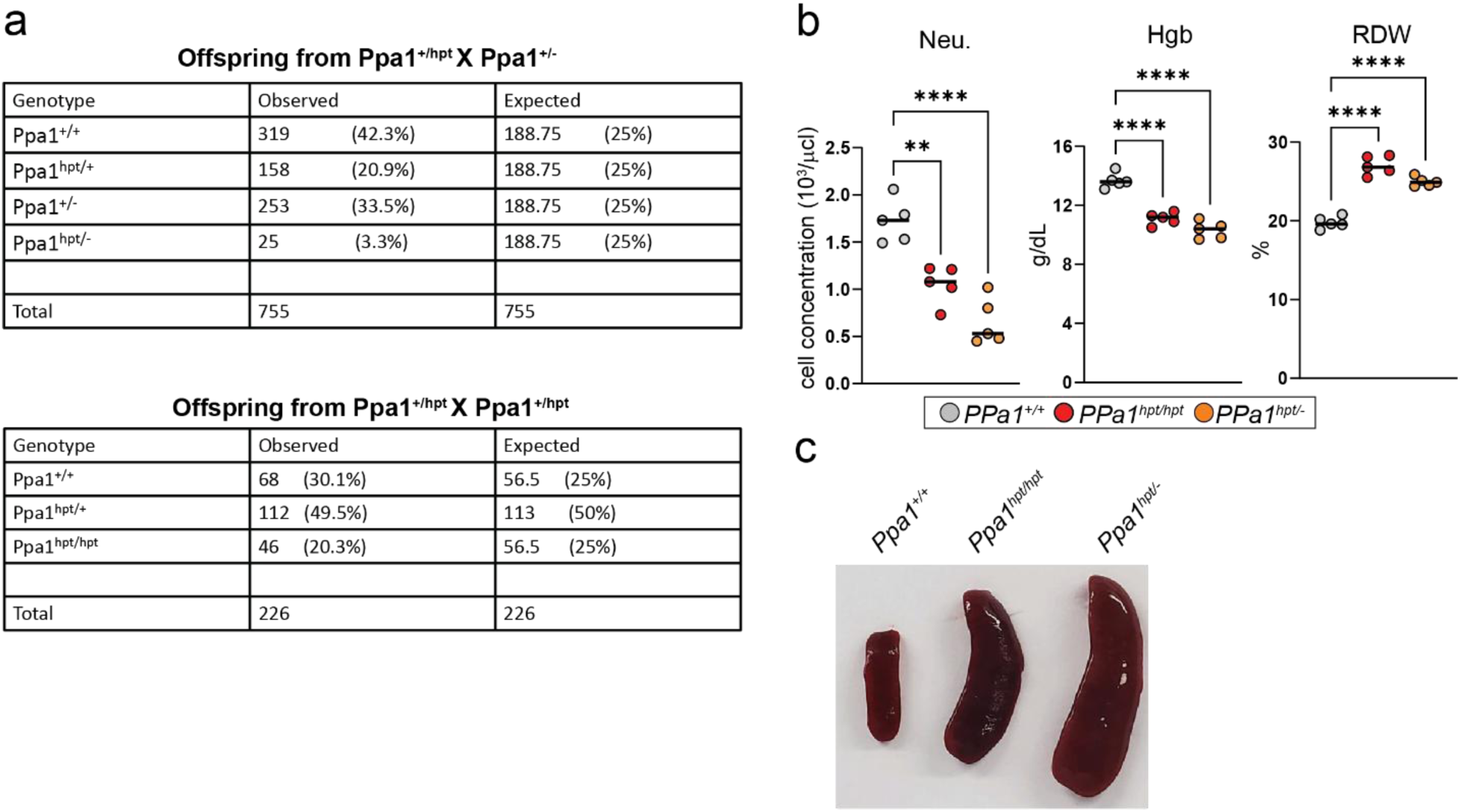
Comparison of *PPa1^hpt/-^* and *PPa1^hpt/hpt^* mouse strains. **(a)** Observed Mendelian inheritance ratios of *PPa1^hpt/-^*and *PPa1^hpt/hpt^* mice from the indicated breeding crosses. **(b)** Peripheral blood neutrophil counts and RBC indices in *PPa1^hpt/-^* and *PPa1^hpt/hpt^* mice. Each symbol represents an individual mouse. Horizontal bars represent the median and statistical significance was determined with one-way ANOVA with Dunnett’s multiple comparisons. **(c)** Representative splenomegaly in *PPa1^hpt/-^* and *PPa1^hpt/hpt^*mice.

**Figure S2:**
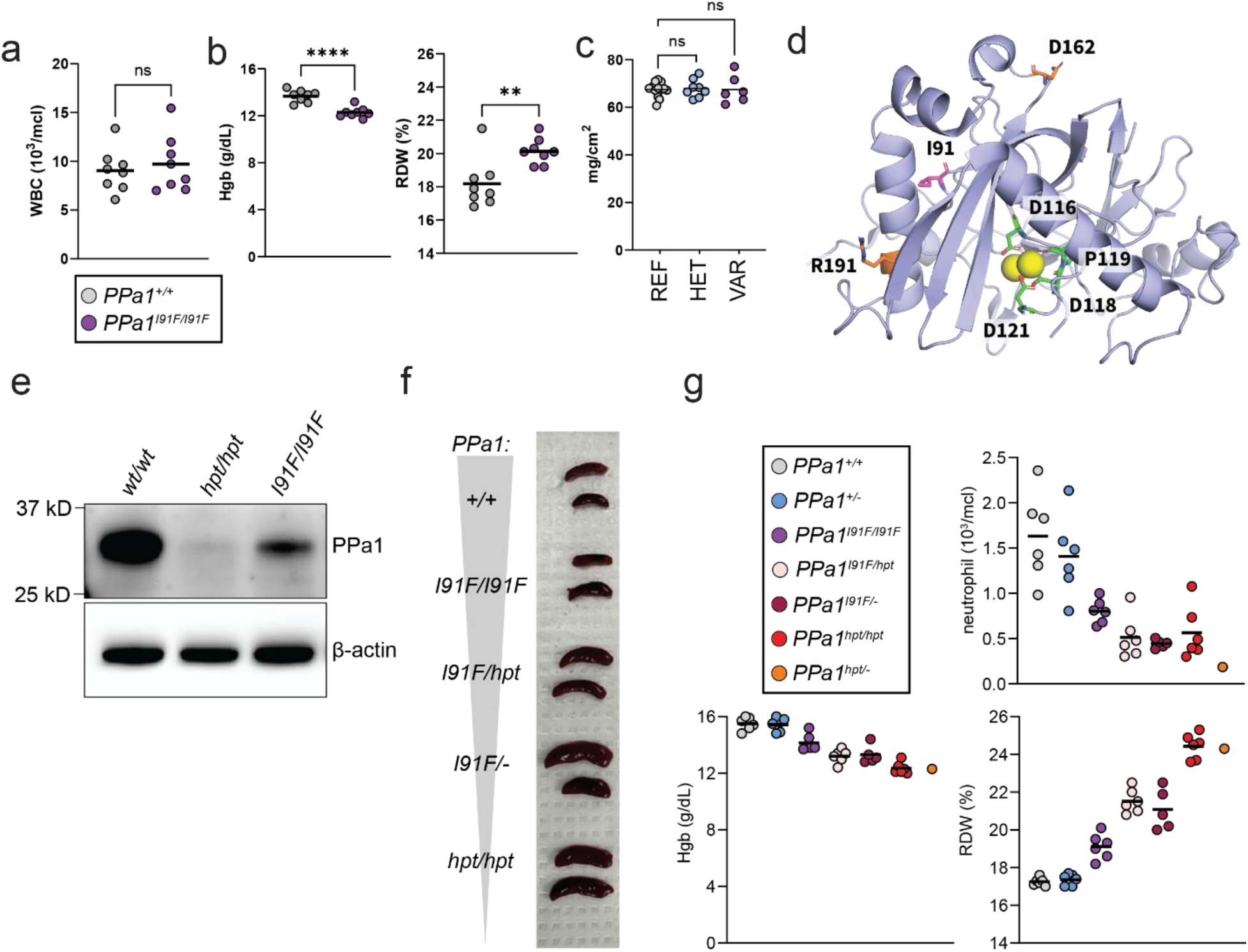
Characterization of PPa1^I91F/I91F^ mutant mice. **(a)** Peripheral WBC counts in *PPa1^I91F/I91F^* mice. **(b)** RBC indices in *PPa1^I91F/I91F^* mice. **(c)** Tibial bone mineral density in *PPa1^I91F/I91F^* mice. **(d)** Structural localization of I91 (purple) within the crystal structure of human PPA1, shown relative to the active site (green) and R191 and D162 (orange). **(e)** Immunoblot analysis of PPa1 expression in bone marrow from the indicated genotypes. **(f)** Spleen size as it relates to *PPa1* gene dosing. **(g)** Peripheral neutrophil counts and RBC indices across an allelic series of PPa1 variants. Each symbol represents an individual mouse. Horizontal bars represent the mean and statistical significance was determined with two-tailed unpaired t-test **(a, b)** and one-way ANOVA with Dunnett’s multiple comparisons **(c)**.

**Figure S3:**
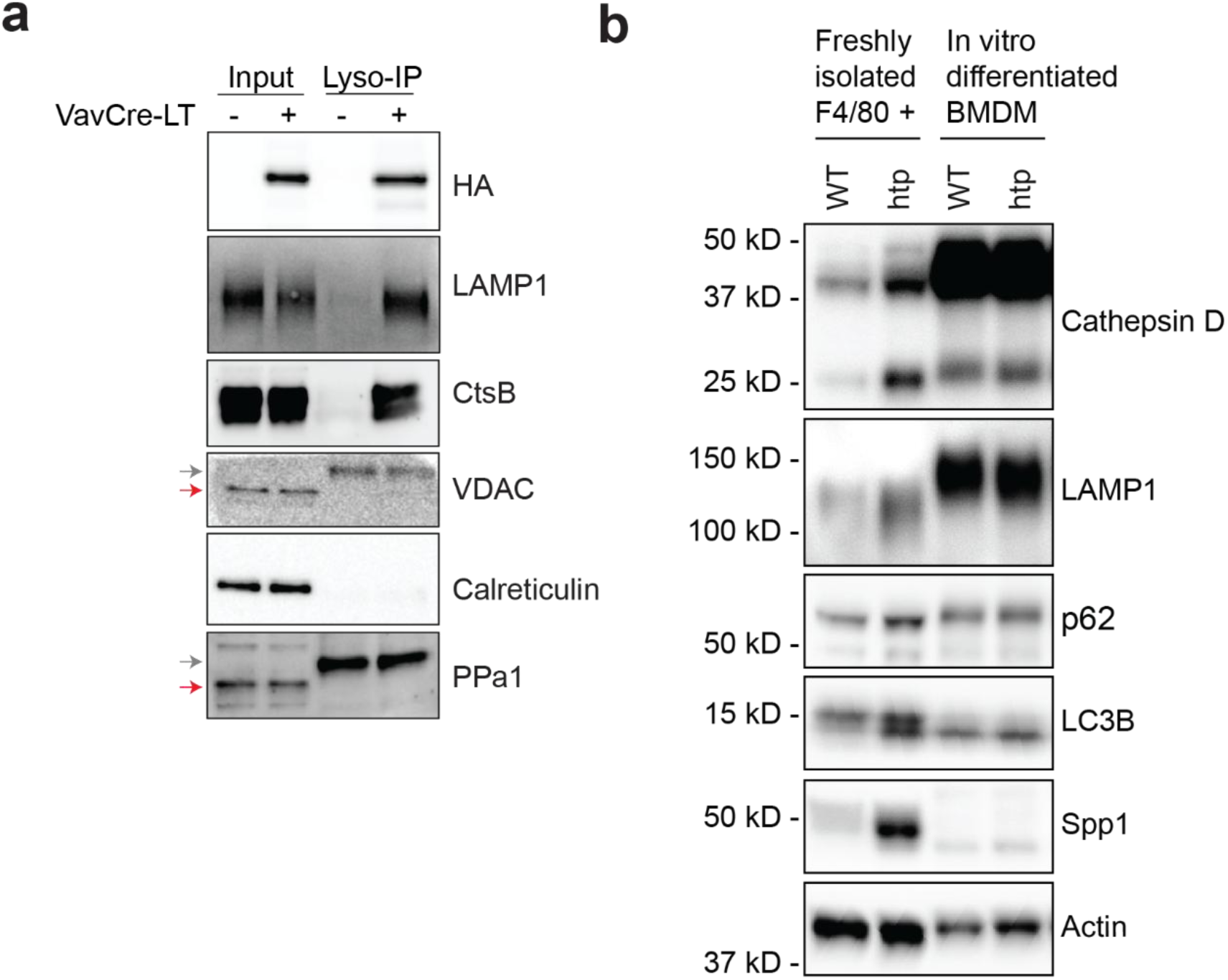
Lysosomal markers in PPa1-deficient cells. **(a)** Lyso-IP from mice expressing 3xHA-TMEM192 within their hematopoietic cells showing that PPa1 does not appear in the lysosomal fraction. Red arrows indicate expected band, gray arrows indicate background band. Data is representative of two independent Lyso-IP experiments. **(b)** Comparison of lysosomal markers in freshly isolated F4/80+ bone marrow macrophages and BMDMs differentiated in vitro for 6 days with CSF-1. Data is representative of three independent BMDM culture experiments.

